## Supplemental Figures for "DNA methylation insulates genic regions from CTCF loops near nuclear speckles"

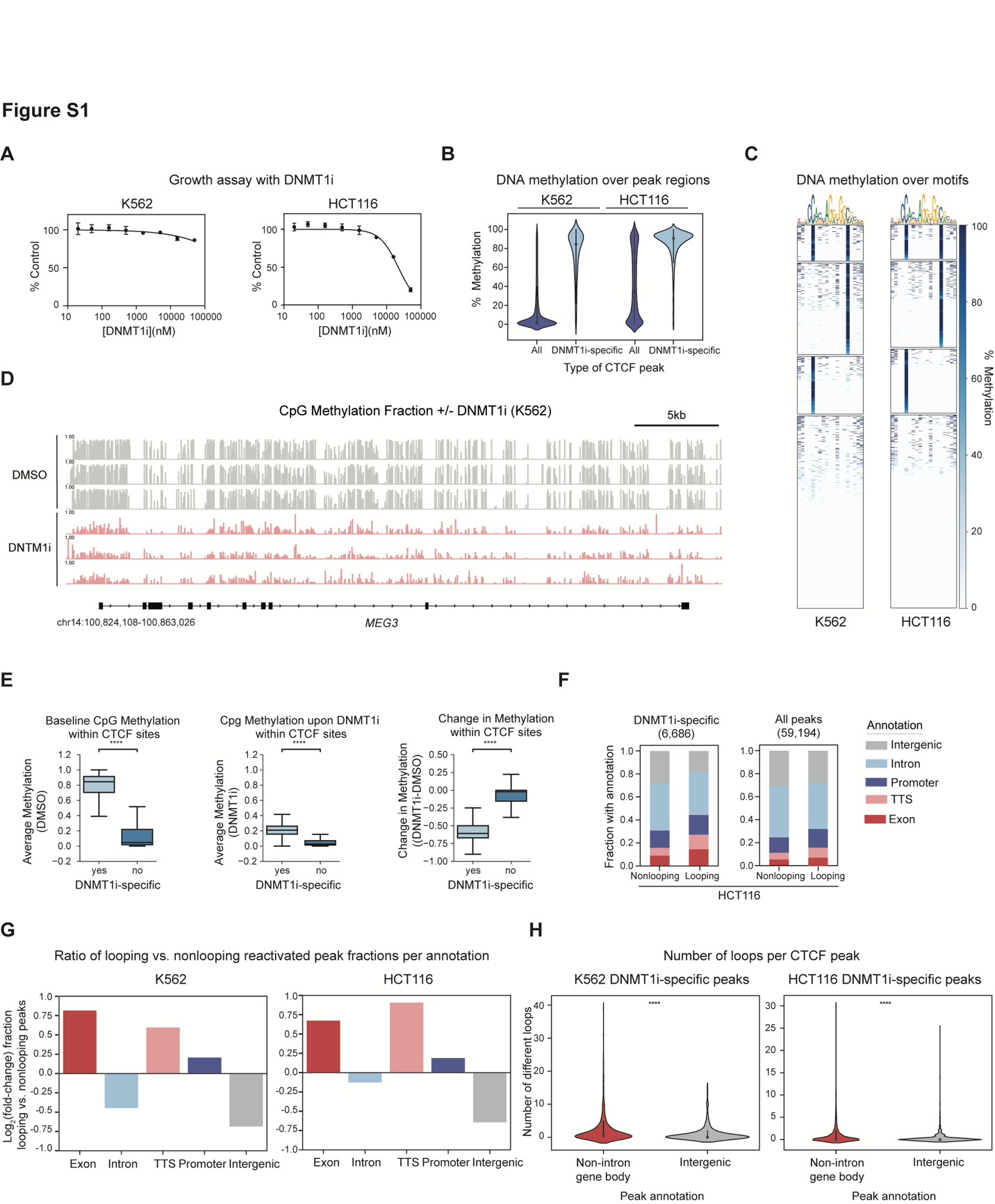
**Figure S1** | DNMT1 inhibition activates CTCF peaks and loops on gene bodies

A. Dose-response curve in K562 (left) and HCT116 cells (right) of percent growth (y-axis) in varying doses of DNMT1i treatment (x-axis) relative to DMSO-treated control cells after 3 days of treatment. Data represent mean ± s.e.m.
B. Percent methylation in wildtype cells across CpG sites within 300 bp CTCF peak regions for DNMT1i-specific peaks and all other CTCF peaks.
C. Percent methylation status of CTCF motifs underlying DNMT1i-specific peaks clustered by K-means. Number of CTCF sites per cluster: K562 (cluster 1: 139, cluster 2: 351, cluster 3: 221, cluster 4: 738); HCT116 (cluster 1: 405, cluster 2: 1005, cluster 3: 746, cluster 4: 2249). Bisulfite data for HCT116 from GEO GSM3317488, K562 from ENCODE ENCFF459XNY.
D. Example Integrated Genome Browser (IGV) CpG methylation fraction ± DNMT1i at the MEG3 locus for K562. K562 CpG methylation data extracted from LIMe-Hi-C methylation bigwigs published in Siegenfeld et. al., 2022.
E. Boxplots showing normalized CpG Methylation fraction (y-axis) averaged across all CpG sites within non-DNMT1i-specific and DNMT1i-specific peaks (x-axis) in the DMSO condition alone (right), the DNMT1i condition alone (middle), and the difference between DNTM1i and DMSO (left). K562 CpG methylation data extracted from LIMe-Hi-C methylation bigwigs published in Siegenfeld et. al., 2022.
F. Proportion of looping and nonlooping peaks with each annotation (y-axis) for both DNMT1i-specific and all CTCF peaks in HCT116 in DNMT1i.
G. log_2_(fold-change) ratio of the fractions of each annotation within looping vs. nonlooping DNMT1i-specific peaks. Ratio of bars from **Fig 1G, S1F**.
H. Number of different loops each CTCF peak makes (number of interaction partners) (y-axis) for DNMT1i-specific K562 (left) and DNMT1i-specific HCT116 (right) peaks in DNMT1i by peak annotation (x-axis).

In (**E**), the interquartile range (IQR) is depicted by the violin with the median represented by the center dot. Outliers are excluded. For **E** and **H**, P values were calculated by a Mann-Whitney test and are annotated as follows: ns: not significant; *: 0.01 <p ≤ 0.05; **: 0.001 <p ≤ 0.01; ***: 0.0001 <p ≤ 0.001; ****: p ≤ 0.0001.

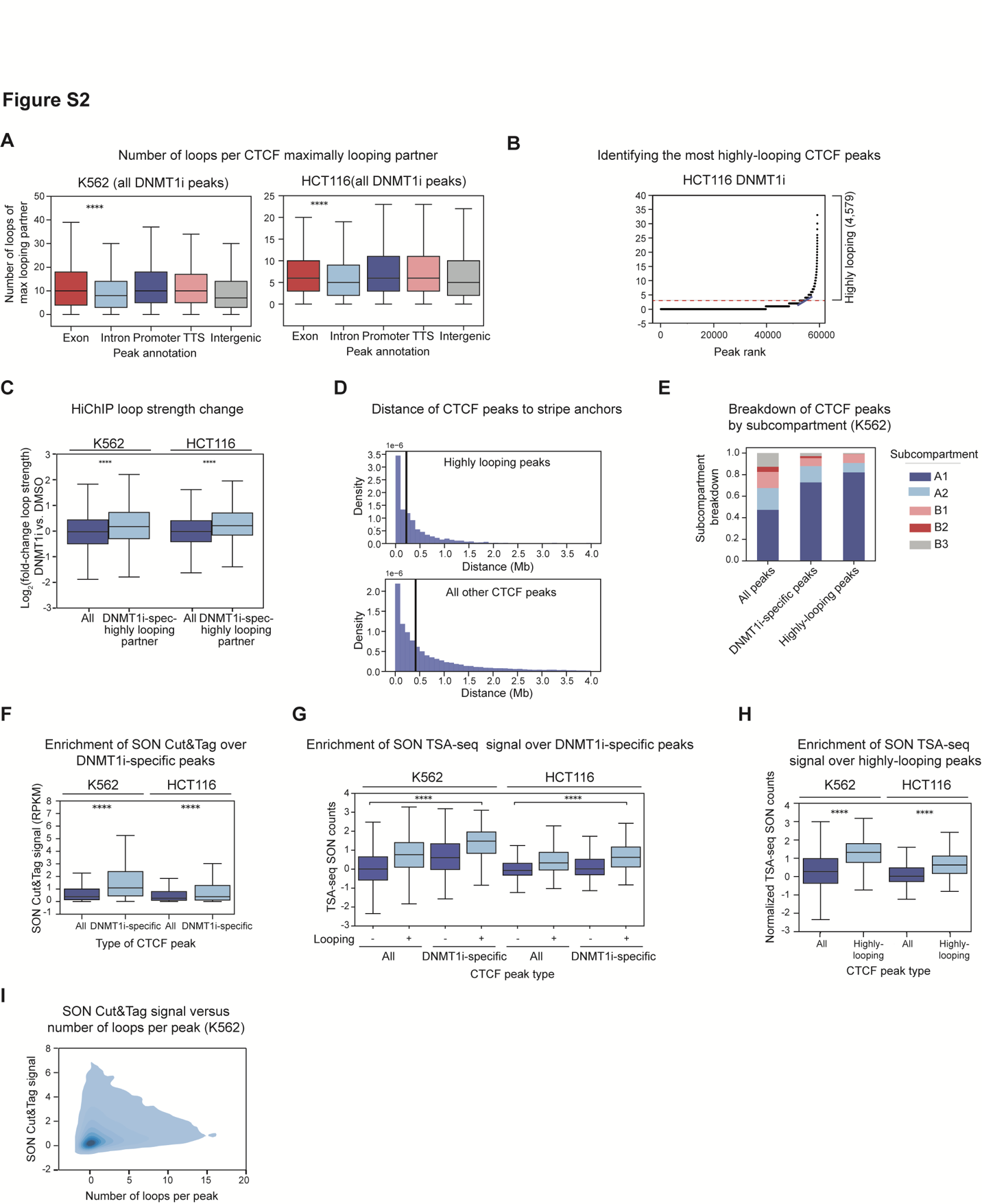

**Figure S2** | DNMT1i-specific CTCF peaks interact with highly-looping partners near nuclear speckles

A. Number of different partner peaks the maximally-looping partner interacts with (y-axis) for all K562 (left) and all HCT116 (right) peaks in DNMT1i by peak annotation (x-axis).
B. Number of different loops (partner peaks) (y-axis) assigned to each peak by rank (x-axis) in DNMT1i for HCT116. Highly-looping peaks reside above the red dashed line.
C. Boxplot showing the log2(fold-change) in loop strength DNMT1i vs. DMSO (y-axis) for loops connecting DNMT1i-specific peaks to highly-looping partners vs. all other loops (x-axis) for K562 and HCT116.
D. Histogram showing density (y-axis) of genomic distances (x-axis) of CTCF highly-looping peaks (top) and all other CTCF peaks (bottom) in DNMT1i-treated K562 cells from stripe anchors identified from DNMT1i-treated K562 Hi-C data. Median distance is shown with a solid black line. K562 LIMe-Hi-C data from Siegenfeld et. al., 2022.
E. Proportion of all, DNMT1i-specific, and highly-looping CTCF K562 DNMT1i peaks in each published subcompartment (y-axis) assigned by SNIPER for wildtype K562 cells.
F. Boxplot showing replicate-averaged SON Cut&Tag signal (RPKM, 20 kb bins, y-axis) after DNMT1i treatment in the respective cell type over DNMT1i-specific peaks vs. all other CTCF peaks called in DNMT1i treatment for K562 and HCT116 cells. Same as **Fig 2G**, but not broken into looping categories.
G. Boxplots showing SON TSA-seq normalized counts (y-axis) for DNMT1i-specific vs. non-DNMT1i-specific CTCF DNMT1i peaks for K562 and HCT116 (left) broken down by whether the CTCF peak is in a loop anchor.
H. Boxplots showing SON TSA-seq normalized counts (y-axis) for highly-looping vs. normal-looping CTCF DNMT1i peaks for K562 and HCT116 (right). SON TSA-seq normalized counts were analyzed from public data (4DNFIVZSO9RI.bw, 4DNFIBY8G6RZ.bw) in the untreated, wildtype respective cell type (see Methods).
I. Density plot showing the correlation between SON Cut&Tag signal (RPKM, 20 kb bins) over a given CTCF peak (y-axis) vs. the number of different loops (partners) assigned to that CTCF peak (x-axis) for K562 DNMT1i treatment.

In (**A**, **C**, **F**, **G**, and **H**), the interquartile range (IQR) is depicted by the box with the median represented by the center line. Outliers are excluded. P values were calculated by a Mann-Whitney test and are annotated as follows: ns: not significant; *: 0.01 <p ≤ 0.05; **: 0.001 <p ≤ 0.01; ***: 0.0001 <p ≤ 0.001; ****: p ≤ 0.0001.

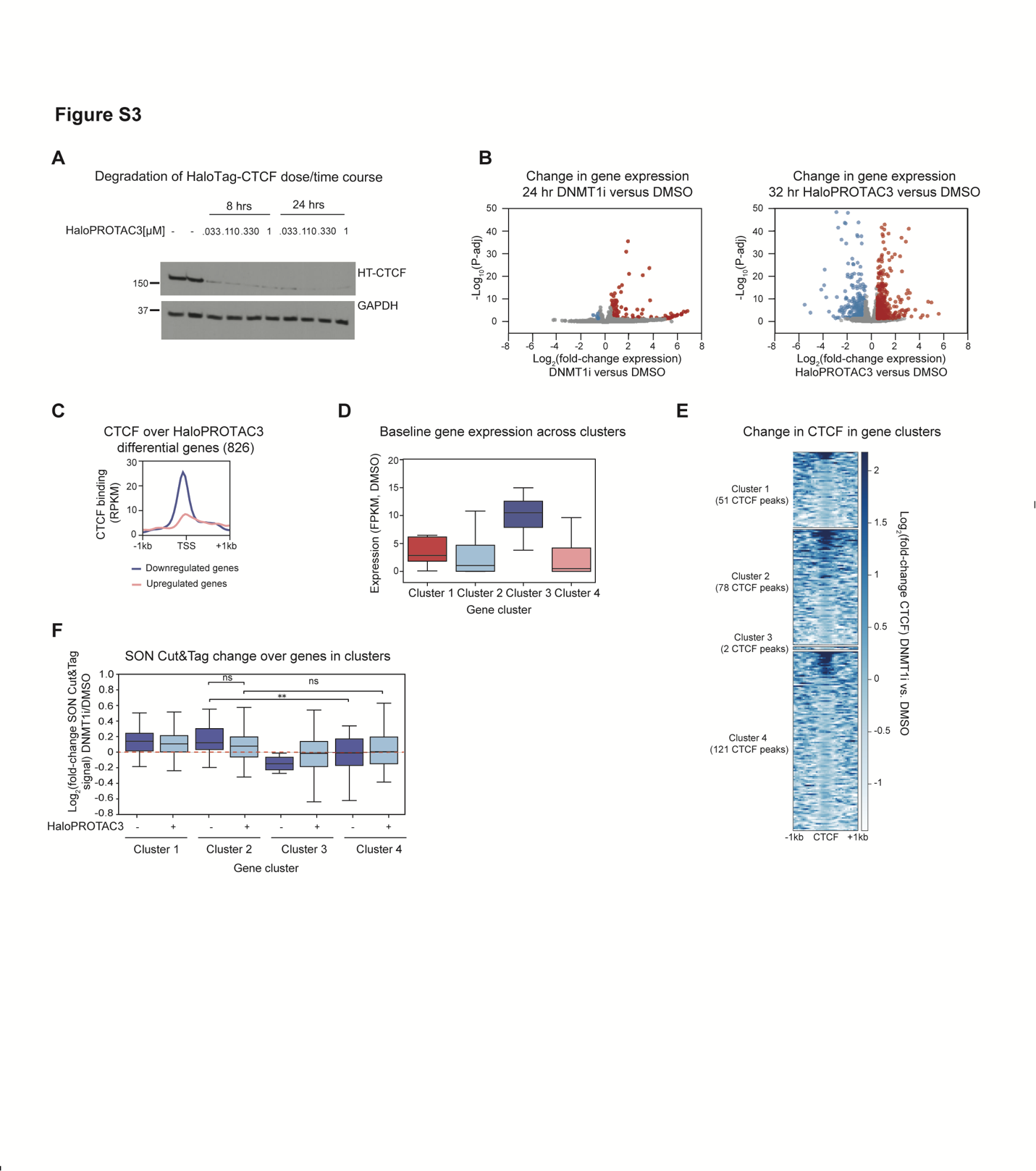
**Figure S3** | DNMT1i-induced gene activation and speckle association depend on CTCF

A. Western blot for CTCF levels over a dose/time course of HaloPROTAC3 in HaloTag-CTCF knock- in HCT116 cells. GAPDH is shown as a loading control. 330 nM was chosen as the final dose.
B. Volcano plot showing -log_10_(P-adj) (y-axis) vs. log_2_(fold-change) in gene expression (x-axis) upon 24 hours DNMT1i treatment (left) or 32 hours HaloPROTAC3 treatment (right) in HaloTag-CTCF HCT116 cells. Upregulated genes are highlighted in red, and downregulated genes are highlighted in blue (P-adj < 0.05, |log_2_(fold-change)| > 0.5). <5 points per plot are omitted for visualization.
C. Aggregate profile plot of replicate-averaged RPKM normalized CTCF binding in DMSO over the transcription start sites of differential HaloPROTAC3 genes.
D. Boxplot showing expression levels of genes within clusters from **Fig 3D** in Fragments Per Kilobase per Million mapped fragments (FPKM). Expression in cells treated with vehicle (- HaloPROTAC3, - DNMT1i) is shown.
E. Heatmap of per-CTCF peak log_2_(fold-change CTCF binding) after 1 day of DNMT1i treatment vs. DMSO control centered over all CTCF peaks overlapping genes in the clusters from **Fig 3D**. Each gene could have multiple CTCF peaks on it.
F. Boxplot showing log_2_(fold-change SON Cut&Tag signal DNMT1i vs. DMSO, y axis) over gene bodies for genes in the different clusters from **Fig 3D**. log_2_(fold-change Cut&Tag signal with and without HaloPROTAC3 treatment are shown for each gene cluster. The same genes are shown for the + and - HaloPROTAC3 conditions for comparison.

In (**D** and **F**), the interquartile range (IQR) is depicted by the box with the median represented by the center line. Outliers are excluded. P values were calculated by a Mann-Whitney test and are annotated as follows: ns: not significant; *: 0.01 <p ≤ 0.05; **: 0.001 <p ≤ 0.01; ***: 0.0001 <p ≤ 0.001; ****: p ≤ 0.0001.

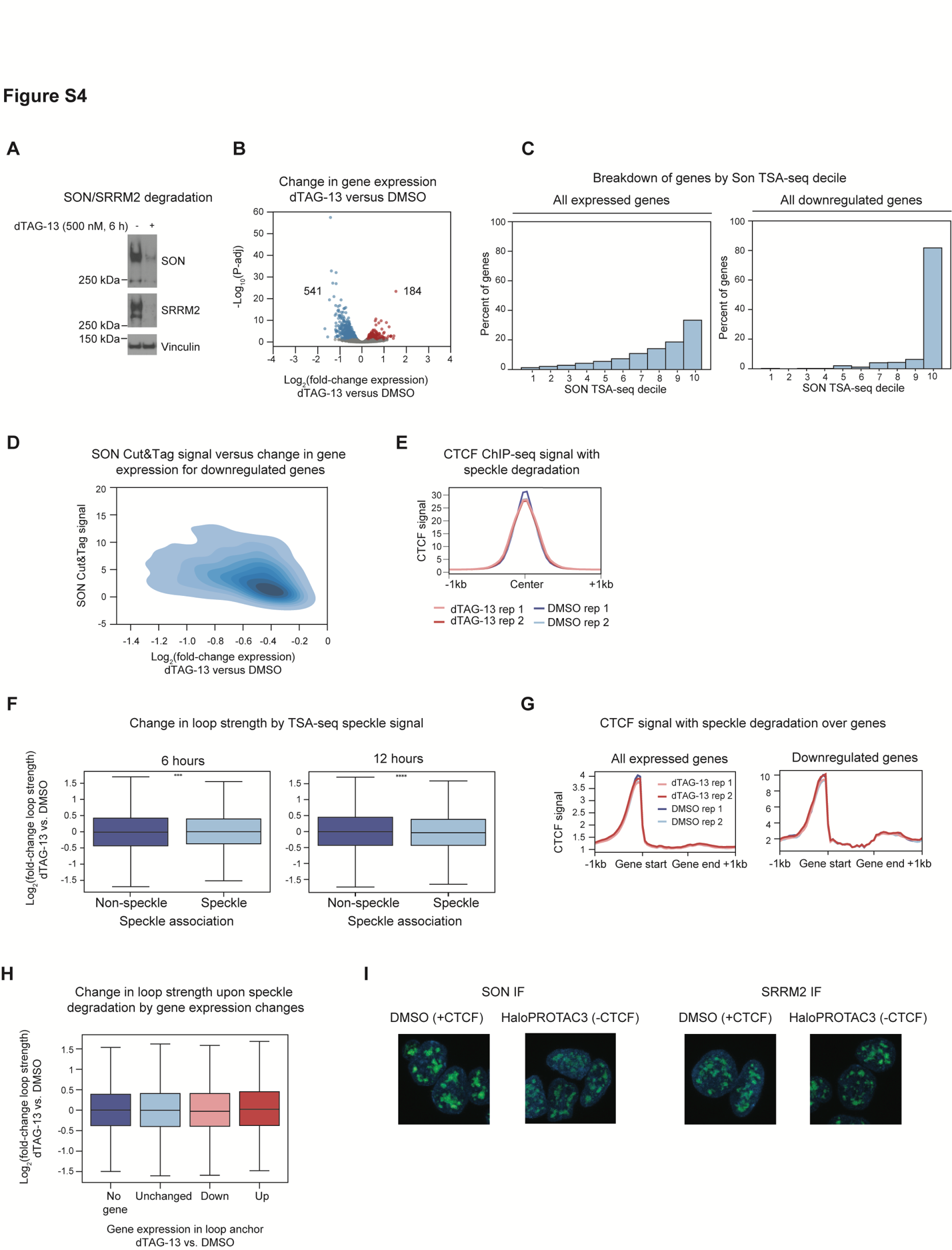

**Figure S4** | Acute disruption of nuclear speckles alters gene expression without disrupting CTCF

A. Separate western blots for SON and SRRM2 in speckle dTAG knock-in K562 cells ± 6 hours of 500 nM dTAG-13 treatment. Vinculin was used as a loading control.
B. Volcano plot showing -log10(P-adj) (y-axis) vs. log2(fold-change) in gene expression (x- axis) upon speckle degradation with 6 hours dTAG-13 treatment. Upregulated genes are highlighted in red, and downregulated genes are highlighted in blue (P-adj<0.05, |log2(fold-change| > 0.25).
C. Proportion (y-axis) of all nonzero (left) and all 6-hour dTAG-13 downregulated (right, P-adj<0.05) genes in each SON TSA-seq decile (x-axis) from wildtype cells. 10 is the decile closest to speckles.
D. Density plot showing SON Cut&Tag signal in wild-type K562 cells treated with DMSO (RPKM 20kb bins, y- axis) vs. the change in gene expression (x-axis) for genes that decrease in expression upon dTAG-13 treatment (Spearman R, -0.43).
E. Aggregate profile plot of CTCF ChIP-seq signal (RPKM) in speckle knock-in K562 cells with and without 6-hour speckle degradation (y-axis) over all CTCF peaks (x-axis).
F. Boxplot showing log2(fold-change) loop strength for dTAG-13 treated vs. DMSO treated speckle dTAG knock-in cells relative to TSA-seq decile after 6 or 12 hours of dTAG-13 treatment with matched DMSO controls. Speckle refers to the decile closest to speckles, and non-speckle refers to the deciles 1-9 that are furthest from speckles.
G. Aggregate profile plot of CTCF ChIP-seq signal (RPKM) in speckle dTAG knock-in K562 cells with dTAG-13 vs. DMSO treatment (y-axis) across expressed genes (top) and genes that are downregulated with dTAG-13 treatment (bottom) (x-axis).
H. Boxplot showing log2(fold-change) loop strength for 6-hour dTAG-13 treated vs. DMSO treated speckle dTAG knock-in cells (y-axis) for loops with anchors that do not overlap any genes, overlap only genes that do not change expression, and overlap genes that increase or decrease in expression for at least one anchor (x-axis).
I. Representative SON and SRRM2 immunofluorescence images (green, alexafluor488) in HaloTag- CTCF HCT116 cells with 24 hours of 330 nM HaloPROTAC3 treatment to deplete CTCF or vehicle control. Overlaid on DAPI in blue.

In (**F** and **H**), the interquartile range (IQR) is depicted by the box with the median represented by the center line. Outliers are excluded. P values were calculated by a Mann-Whitney test and are annotated as follows: ns: not significant; *: 0.01 <p ≤ 0.05; **: 0.001 <p ≤ 0.01; ***: 0.0001 <p ≤ 0.001; ****: p ≤ 0.0001.
